# Disentangling the dynamics of energy allocation to provide a proxy of robustness in fattening pigs

**DOI:** 10.1101/2022.10.19.512827

**Authors:** Guillaume Lenoir, Loïc Flatres-Grall, Rafael Muñoz-Tamayo, Ingrid David, Nicolas C. Friggens

**Author notes:** E-mail addresses: GL LFG RMT ID NF.

## Abstract

**Background:** There is a growing need to improve robustness characteristics in fattening pigs, but this trait is difficult to phenotype. Our first objective was to develop a robustness proxy on the basis of modelling of longitudinal energetic allocation coefficient to growth for fattening pigs. Consequently, the environmental variance of this allocation coefficient was considered as a proxy of robustness. The second objective was to estimate its genetic parameters and correlation with traits under selection as well with phenotypes routinely collected on farms. A total of 5848 pigs, from Piétrain NN paternal line, were tested at the AXIOM boar testing station (Azay-sur-Indre, France) from 2015 to 2022. This farm was equipped with automatic feeding system, recording individual weight and feed intake at each visit. We used a dynamic linear regression model to characterize the evolution of the allocation coefficient between cumulative net energy available, estimated from feed intake, and cumulative weight gain during fattening period. Longitudinal energetic allocation coefficients were analysed using a two-step approach, to estimate both its genetic variance and the genetic variance in the residual variance, trait LSR.

**Results:** The LSR trait, that could be interpreted as an indicator of the response of the animal to perturbations/stress, showed low heritability (0.05±0.01). The trait LSR had high favourable genetic correlations with average daily growth (−0.71±0.06) and unfavourable with feed conversion ratio (−0.76±0.06) and residual feed intake (−0.83±0.06). The analysis of the relationship between estimated breeding values (EBV) LSR quartiles and phenotypes routinely collected on farms shows the most favourable situation for animals from quartile with the weakest EBV LSR, *i.e*., the most robust.

**Conclusions:** These results show that selection for robustness based on deviation from energetic allocation coefficient to growth can be considered in breeding programs for fattening pigs.

## Background

The pig industry faces new challenges related to rapidly changing environmental conditions, especially related to global warming (Hansen et al., 2012), and to growing societal concerns. For several decades, breeding objectives were mainly focused on increasing animal productivity (growth, feed efficiency…) at the expense of non-productive functions, *i.e*., fitness (Puillet et al., 2016; Rauw et al., 1998). These unfavorable consequences could be explained by trade-offs in resource allocation between biological functions (Rauw, 2009). Indeed, when animals cannot obtain more resources, *i.e*., in limiting environments, allocation of these resources to a high priority function must be to the detriment of another function (Stearns, 1992). In this situation the animal is unable to maximize the expression of each biological function simultaneously. This changing context requires having animals able to adapt to new environmental conditions with more limiting resources, which can be associated with an improvement of robustness. Knap (2005) defined robustness as “*the ability to combine a high production potential with resilience to stressors, allowing for unproblematic expression of a high production potential in a wide variety of environmental conditions”*. Generally, the production potential is associated to a phenotype of interest, such as growth, feed efficiency or milk production, egg production. Incorporating one or several traits to evaluate robustness of growing pigs in genetic evaluation would therefore be of value for the development of more sustainable breeding goals (Berghof et al., 2019b). Accordingly, when targeting robustness as a breeding objective, it is important to maintain simultaneously a high level of production to meet industry’s economic expectations. However, until recently to it was very difficult to phenotype traits such as robustness in farm animals and therefore on the way to improve it.

In parallel, the increasingly common use of sensors in farms, especially automatic feeding system (AFS) in pig industry, allows continuous individual recording of weight or feed intake over a long period. This offers the possibility to characterize the dynamics of those phenotypes for each individual in the face of variations in the environment. Several studies have used such longitudinal data to quantify resilience indicators based on deviation between an expected trajectory of each individual for a given non-perturbed environment and its observed trajectory on feed intake (Nguyen-Ba et al., 2020) or body weight (Revilla et al., 2022). Definition and modelling of individual potential trajectory are challenging issues in these approaches. Other studies developped several resilience indicators based on the within individual variance of time series mesurments related to production, such as feed intake of growing pigs (Putz et al., 2019), milk yield for dairy cows (Poppe et al., 2021) or egg production in laying hens (Bedere et al., 2022). These modeling approaches have mainly addressed the characterization of robustness or resilience through the analysis of one production variable. They are a substantial contribution in the phenotyping of resilience but do not address the underlying biological mechanisms and the potential trade-offs in the use of available resources between production and other functions. A robust animal can be considered as an animal able to allocate a proportion of its resources to the right function at the right time (Friggens et al., 2017). To our knowledge, the characterization of robustness based on the temporal evolution of the allocation pattern has been little explored in growing pigs.

The acquisition of temporal data of feed intake and weight in growing pigs made it possible to consider the development of allocation model based on these two variables to caracterize robustness. With this objective, we developed a conceptual model to represent the temporal pattern of allocation of energy intake to growth in fattening pigs (Lenoir et al., 2022).

In the present study, the first objective was to develop a robustness indicator on the basis of modelling of longitudinal energetic allocation coefficient to growth for fattening pigs. Consequently, the environmental variance of this allocation coefficient was considered as a proxy of robustness. This proxy should reflect the ability of an animal to express or adapt its production potential in the face of changes in the environment relative to other animals that have been raised under the same conditions. Our objective was to estimate its genetic parameters and correlation with traits under selection as well as with phenotypes routinely collected on farms and associated to robustness or heatlh status.

## Methods

### Study population

A total of 25745 pigs from Piétrain NN Français paternal line (Pie NN), free from halothane-sensitivity, of the AXIOM company were used in this study. Individuals from the Pie NN line were born in 3 different farms integrated into the AXIOM breeding scheme and that comply with AXIOM’s biosafety and health requirements. A part of the males were selected before weaning and then raised at the boar testing station of the breeding company AXIOM Genetics (Azay-sur-Indre, France). The animals considered in the present dataset were 6885 entire males and 13012 females raised and individually tested at their farm of birth from April 2014 to April 2022 and 5848 entire males raised from September 2015 to April 2022 at the boar testing station.

The animals raised on their birth farm were born from 3943 litters, 6.5±2.9 piglets per litter, and from 321 sires, 80±53.8 piglets per sire. To limit the risk of confounding between environmental (*i.e*., fattening group) and genetic effects, the sires were used at least in two mating groups in each farm and in two different farms. Animals were transferred to fattening rooms when they were 75.7 ±3.4 days of age (33.8±7.8 kg body weight (BW)). Pigs were raised in fattening rooms for 68.6 ±4.9 days until the individual testing at around 142.4±4.6 days of age (103.4±11 kg BW).

For males raised at the boar testing station, they were transferred every three weeks from birth farm to the station at an average age of 27.3±2.2 days with an average BW of 8.5±1.7 kg. They were raised in pens of 14 animals from the same birth farm. These groups of 14 pigs were never modified at the different stages of breeding. Each fattening group consisted of animals sourced from between one or three farrowing farms. These animals came from 2048 litters, 2.6±1.5 piglets per litter, and were born from 238 sires, 22.1 ±15 piglets per sire. They were raised in quarantine and post-weaning rooms for five and two weeks respectively and transferred to fattening rooms when they were 76.4±2.9 days of age (34.4±5.4 kg BW). These pigs were raised in fattening rooms for 69 ±4.7 days until the individual testing at around 145.4 ±3.6 days of age (104.5±11.1 kg BW). Fattening rooms were equipped with AFS: Nedap pig performance testing feeding station (Nedap N.V.; Groenlo, the Netherlands). Animals were fed ad-libitum with commercial diets adapted to their physiological needs. The provided diets were non-limiting in amino acids. The boar testing station environmental and technical conditions are described in detail in Lenoir et al. (2022a). The pedigrees contained 27276 animals across 20 generations.

### Information recorded during the fattening period

The performances recorded were the same in farrowing farms testing and boar testing station. Each animal was individually weighed on arrival in the fattening room (initial body weight: IBW). When the average weight of the group was approximately 100 kg, individual tests were performed for animals weighing more than 70 kg (Institut Technique du Porc, 2004). Measurements made during the test were: body weight (TBW), average ultrasonic backfat thickness (BF = mean of three measurements in mm) and ultrasonic longissimus dorsi thickness (LD = one measurement in mm). The BF and LD measures were transformed to correspond to their values at 100 kg liveweight (BF100 and LD100 respectively) to compare animals at equivalent weight. This transformation was done by applying linear coefficients that multiply the difference between 100 kg and TBW. Coefficients used are 0.04 mm/kg for BF100 and 0.27 mm/kg for LD100 (Sourdioux et al., 2009). The average daily gain (ADG) was calculated as the difference between TBW and IBW divided by the number of days elapsed between the two weighings.

Additionally, at the boar testing station, BW (kg) and feed intake (FI; kg per visit) were recorded each time the animal went into the AFS. The feed conversion ratio (FCR) was calculated as the ratio between the total FI during the fattening period and the weight gain (TBW-IBW), expressed in kg/kg. The average daily feed intake (DFI) was calculated as the total FI during the period divided by the number of days elapsed. The residual feed intake (RFI) was also estimated for each animal as the deviation between the recorded DFI and the potential average daily feed intake (PDFI) predicted from requirements for maintenance and production. Based on the method proposed by Labroue et al. (1999), the PDFI was estimated by linear regression, with the lm function in R (R Core Team, 2018), of DFI on average metabolic weight (AMW), ADG and BF100. The AMW was estimated for each animal using the formula proposed by Noblet et al. (1991), 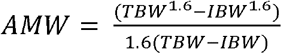. In addition, all medical treatments received by the animal were recorded. A visual observation of the animals was carried out by the technician in charge of the measurements in order to note any morphological defects, anomalies and clinical signs of disease according to a frame of reference (Institut Technique du Porc, 2004), noted as “observable defects”. These observations were made by the same person within any given fattening group. The medical treatments and individual observations were recorded from January 2019 to April 2022 on 3028 males fattened at the boar testing station.

### Longitudinal data pre-treatment

A pre-treatment process was performed on BW and FI, recorded each time the animal went into the AFS, to validate them, identify quality issues and convert them on a daily scale. This process followed the procedure proposed by Revilla et al. (2022) and modified by Lenoir et al. (2022a). In summary, on the scale of the visit, a quadratic regression of BW on age + age^2^ for each animal was applied to eliminate aberrant BW. For a given animal and a given visit, if the ratio between the residual value and the fitted value was > 0.15, the BW measurement was considered to be null. This step was repeated a second time. Following this step, the body weight (BW_*it*_; kg) was estimated from the median of the non-null weights for each pig (*i*) and each fattening day since the transfer to fattening room (*t*). For feed intake, if for a given animal the feed intake rate at a visit was lower or higher than its mean take rate over the fattening period ±4 standard deviations, the FI measurement was considered to be missing. This missing value was estimated using a linear regression of FI on feeding duration. The daily feed intake (FI_*it*_; kg) was calculated as the intakes during the visits of the day *t*. Then BW_*it*_ and FI_*it*_ were validated at the pen scale to detect inconsistencies linked to the AFS machine. When a control day was missing (due to a mechanic problem of AFS or loss of a RFID tag), the missing BW_*it*_ and FI_*it*_ were estimated separately by using local regression model, “*proc loess*” implement in SAS (SAS Institute Inc., 2013). Data recorded on day *t*=0 were excluded from the dataset due to AFS calibration and animal adaptation. After data pre-treatment, the file included 405983 daily records associated to the 5848 males fattened at the testing station.

### Model for analysis

#### Modelling energetic allocation coefficient to growth

As shown on Figure 1, the feed intake, i.e input of the system, is transformed in net energy intake and allocated to several functions: maintenance, body development (protein deposition), body reserves (lipid deposition) and other functions (van Milgen et al., 2005). The body weight gain, *i.e*., output of the system, is directly related to the protein and lipid depostion. Resource allocation is regulated during the fattening period for each individual according to deterministic factors: genetic potential and degree of maturity (Lewis and Emmans, 2020). Over the time, resource allocation coefficient is also impacted by changes in environmental conditions, *i.e*., pertubations (Friggens et al., 2017).

**Figure 1.**
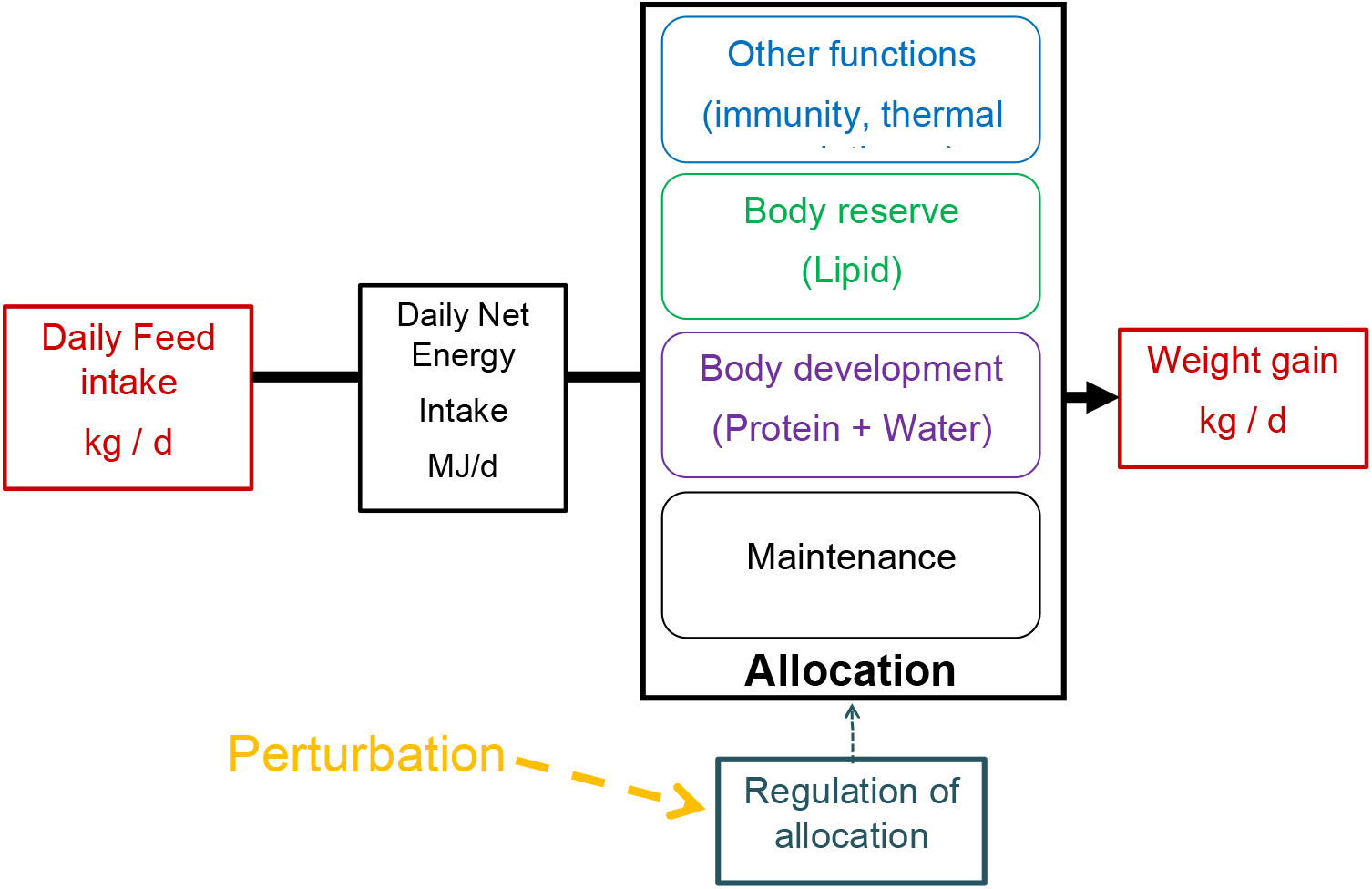
Conceptual model of resource allocation in growing pig. In red : Variables recorded by AFS

A dynamic regression model (DLM; West and Harrison, 1997) was used to estimate daily energetic allocation coefficient to growth (α_*it*_; Lenoir et al., 2022b). First, FI_*it*_ was converted in net energy intake in MJ (EI_*it*_), using the net energy density of the feed of 9.85 MJ of NE/kg. Then, the net energy available for growth at day *t* (NEA_*it*_) was calculated as the difference between EI_*it*_ and the net energy maintenance requirements at day *t* (MR_*it*_), estimated according to Noblet et al. (2016). The DLM to estimate the allocation coefficient of energy to weight gain for a given pig *i* at day *t* (α_*it*_) was built with two equations : an observation equation (1), relating cumulative weight gain at day *t* (CW_*it*_ in kg) and cumulative net energy available at day *t-1* (CNEA_*it-1*_ in MJ), a system equation (2), describing the changes in α_*it*_ (unobserved state variable) from day to day according to a stochastic process.

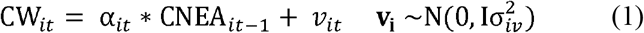

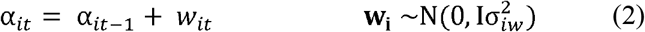

Where *v_it_* was a random observation error for animal *i*; 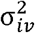 is the observational variance *i*; *w_it_* represented random and unpredictable changes in level between time *t−1* and *t*; and 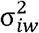 was the system variance. The model was built using the R package dlm (Petris et al., 2009). The values of α_*it*_ were calculated independently for each animal with a Kalman smoother algorithm. The value of α_*it*_ at *t*=1 was not estimated because the consumption at *t-1* was unknown.

##### Estimation of genetic variance in environmental variance

Longitudinal energetic allocation coefficients (α_*it*_) were analyzed with ASReml 4.2 software (Gilmour et al., 2009) to estimate both its genetic variance and the genetic variance in the residual variance (*i.e*., environmental variance) using a two-step approach (SanCristobal-Gaudy et al., 1998; Garreau et al., 2008).

##### First step: estimation of genetic variance in the energetic allocation coefficient

The energetic allocation coefficient was analyzed by a random regression model (RR) with first order Legendre polynomials (Robson, 1959) for the genetic and permanent environmental effects. The common litter was significant as a random effect, tested using likelihood ratio (LRT) test, a-risk of 5%, and included in the model, in addition to additive genetic and permanent environmental effects. Fixed effects included in the model were selected at an p-value of 5% using the Wald F statistic. The significant fixed effects were the fattening group (103 levels), as contemporary group, and the joint effect of fattening group and the fattening pen (517 levels). The age *k* in days of the animal at day *t* was include as a covariate. The residual variance was assumed constant over time.

##### Second step: estimation of genetic variance in residual variance

In the second step of the analysis, the residuals (*e_it_*, residuals of animal *i* at time *t*) of the RR model were used to compute log transformed squared residuals: 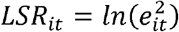 as an indicator of animal robustness. A lower LSR value is assumed to be an indication of a higher animal robustness to environmental perturbations, related to a smaller deviation from expected allocation of energy to growth. To follow the assumption of the BLUP (best linear unbiased prediction; Henderson, 1977) method, which should be applied to a non-selected base population, and to estimate the covariance between traits, a multi-traits animal model including the four traits under selection (ADG, BF100, LD100 and FCR, single measurement for all the animals) and the non-selected traits LSR (repeated data for animal in station) and RFI was applied. For the LSR trait, the same fixed effects were fitted as for α_***it***_ and the random effects included were common litter, permanent environmental and animal additive genetic effects. For the four traits under selection, the fixed effects tested at an α-risk of 5% using the Wald F statistic were the gender (2 levels), the fattening farm (4 levels) and the fattening group within the fattening farm (443 levels). The significant random effects were common litter and animal additive genetic. At this step, heritability (h^2^) was calculated as the ratio of animal genetic variance to the total phenotypic variance, *i.e*., the sum of the genetic additive variance, environmental variances (common litter, permanent environmental if necessary) and the residual variance.

##### Relation between LSR and routinely collected phenotypes

To evaluate whether the LSR phenotype could be considered as a robustness proxy, the relationship between estimated breeding value (EBV) for LSR and health phenotypes was studied. The 3028 males with LSR phenotype and known information over the fattening period (observations, injections…) were divided into 4 quartiles according to their EBV for LSR, from Q1 for the most favorable values (lower EBV LSR) to Q4 for the most unfavorable values (higher EBV LSR). We studied the distribution of other phenotypes associated with animal health and robustness according to the EBV LSR quartile. To compare the differences and frequencies in the scores among the four EBV LSR classes, a Chi-square was performed. Statistical significance was set a priori at P less than or equal to 0.05. These phenotypes are derived from measurements made during the animals performance evaluations, and from the medical treatments recorded during the testing period. In each class, we differentiated animals that can be selected (Selectable) from those that are dead or weighing less than 70 kg at the day of the individual test or weighing 70 kg or more and with an observable defect on the day of testing. We considered as an observable defect on the day of testing, factors such as weak development and similar that were estimated to relate to the robustness of the animal (Appendix 1). A second trait differentiated pigs that received at least one antibiotic or anti-inflammatory injection during the testing period from those that didn’t receive any injection (No injection). We also differentiated pigs that were “Selectable” without receiving any antibiotic or anti-inflammatory injection during the testing period (Selectable without injection) from the others.

## Results

### Observed allocation coefficients and robustness indicators

The descriptive statistics for the dataset used in this study are shown in Table 1. The observed means and LSR were 0.099±0.027 kg/MJ NE and −12.62±2.50, respectively. The phenotypic correlations, estimated with *cor.test* function on R (R Core Team, 2018), for trait α_*t*_ were positive with *e_t_* (0.241±0.002) and LSR (0.23±0.002), which means that a higher energetic allocation rate to growth was related to a higher variability. The phenotypic coefficients of variation were greater than 20% for IBW, α_*t*_ and LSR, and between 10 and 20% for TBW, ADG, DFI and BF100, indicating large phenotypic variation for these traits.

**Table 1.** Descriptive statistics of the variables recorded or estimated on fattening pigs

| Trait (unit)<sup>1</sup> | Number of animals /<br>records if repeated<br>measures | Mean | SD |
| --- | --- | --- | --- |
| **IBW (kg)** | 25745 | 33.8 | 7.8 |
| **TBW (kg)** | 25365 | 103.7 | 11.1 |
| **ADG (kg/d)** | 25322 | 0.977 | 0.109 |
| **FCR (kg/kg)** | 8675 | 2.25 | 0.21 |
| **DFI (kg/d)** | 8675 | 2.19 | 0.29 |
| **RFI (kg/d)** | 8675 | -0.005 | 0.169 |
| **BF100 (mm)** | 25323 | 7.66 | 1.19 |
| **LD100 (mm)** | 25320 | 68.26 | 6.34 |
| **$\alpha_t$ (kg/MJ)** | 5848 / 405104 | 0.099 | 0.027 |
| **$e_t$ (kg/MJ)** | 5848 / 405104 | 0 | 0.0096 |
| **LSR** | 5848 / 405104 | -12.62 | 2.50 |

<sup>1</sup>IBW: initial body weight; TBW: terminal body weight; ADG: average daily gain; FCR: feed conversion; DFI: daily feed intake; RFI: residual feed intake; BF100: backfat thickness estimated at 100kg liveweight; LD100: longissimus dorsi thickness estimated at 100 kg liveweight; $\alpha_t$ : allocation coefficient to growth; $e_t$ : residual of RR model; LSR= log-squared residual, robustness indicator.

Figure 2 displays the α_*t*_ trajectories of two animals exhibiting different patterns. The first animal on Figure 2a had a smoothed allocation trajectory over time close to its prediction from RR model (Figure 2a), its average LSR value for was −14.6±1.7. The second animal on Figure 2b had higher deviation between smoothed allocation and prediction likely in response to an environmental perturbation, its average LSR value was higher than for the first individual (−12.3±1.9). Accordingly, the parameter LSR looks to be a useful indicator to quantify the effect of perturbation of an animal and allows comparison within a population.

**Figure 2.**
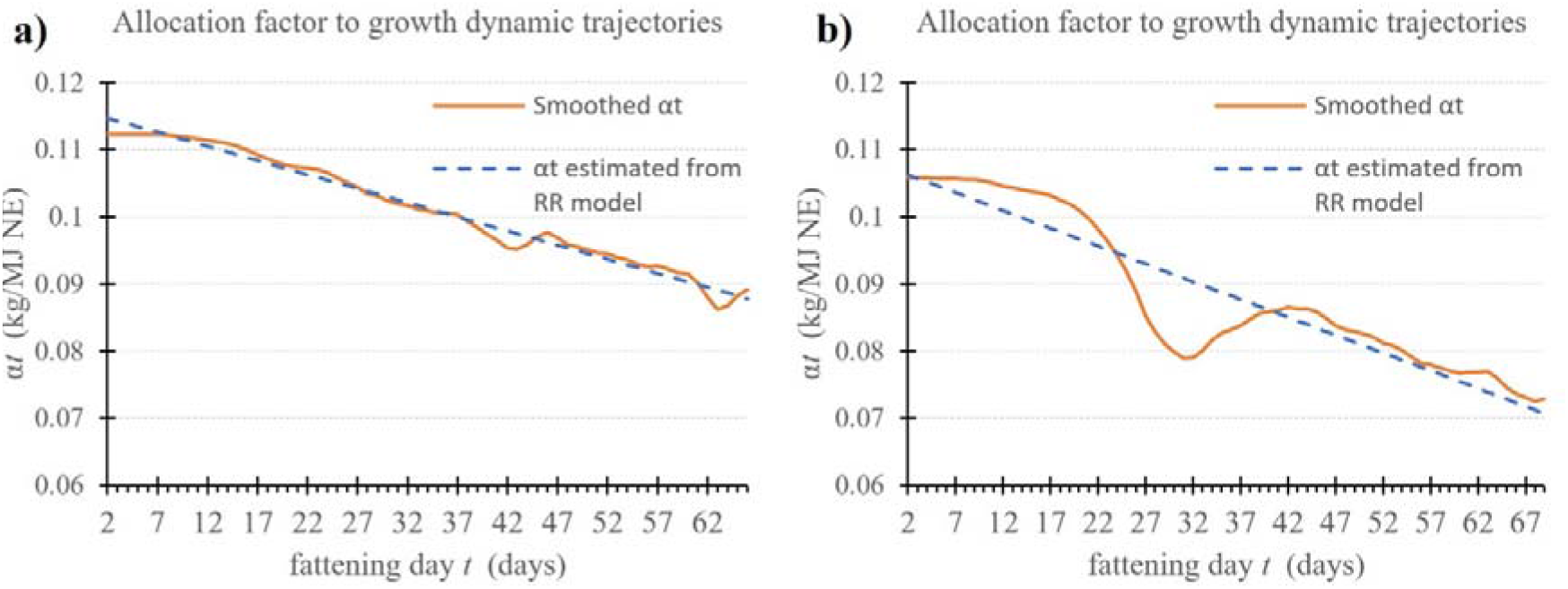
Example of two dynamic trajectories of the allocation coefficients α_*t*_ during the whole fattening period for two animals: smoothed with DLM model (orange line) and its prediction from RR model (blue dotted line).

### Genetic parameters of allocation coefficients, production and robustness indicator traits

The changes in heritability for α_*t*_ over time estimated with the RR model are shown in Figure 3, ranging from 0.20±0.03 to 0.30±0.03. The heritabilities obtained with the RR model were stable from 67 to 100 days of age, around 0.30±0.03, then decreased up to 150 days of age and then stabilized around 0.20±0.03 toward the end of the control period. The permanent environmental ratios ranged from 0.51±0.03 to 0.64±0.03. The estimates obtained decreased up to 128 days of age and then increased again toward the end of the period.

**Figure 3.**
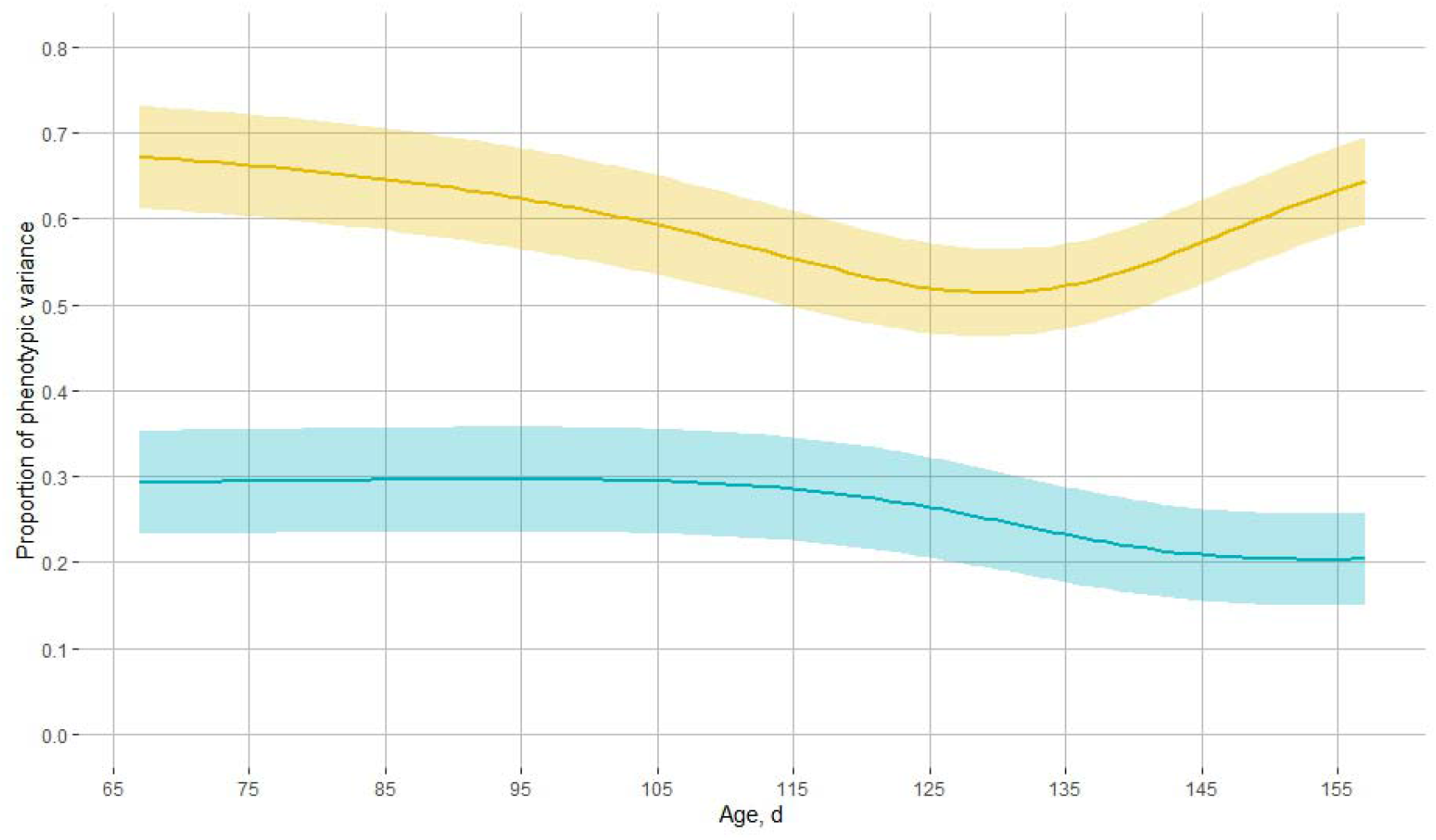
Changes of heritability (*h*^2^_k_; blue) and permanent environmental (*p*^2^_k_; yellow) estimates for energetic allocation coefficient α_*t*_ over age in days under the random regression model (RR) using Legendre orthogonal polynomials. Shaded area: 95% confidence interval

Heritability estimates of the traits under selection, ADG, BF100, LD100 and FCR, were moderate, ranging from 0.27±0.03 to 0.45±0.02 (Table 2). Heritability estimates for RFI and FCR were not significantly different from each other, 0.29±0.03 and 0.27±0.03 respectively. The robustness indicator LSR was lowly heritable, 0.05±0.01. The proportion of variance due to common litter effects was similar for all traits, ranging from 0.04±0.01 to 0.06±0.01, except in the LSR estimation, which had a proportion of phenotypic variance explained by litter effect close to 0. The proportion of phenotypic variance explained by permanent environment effect for LSR was moderate, 0.22±0.01.

**Table 2.**
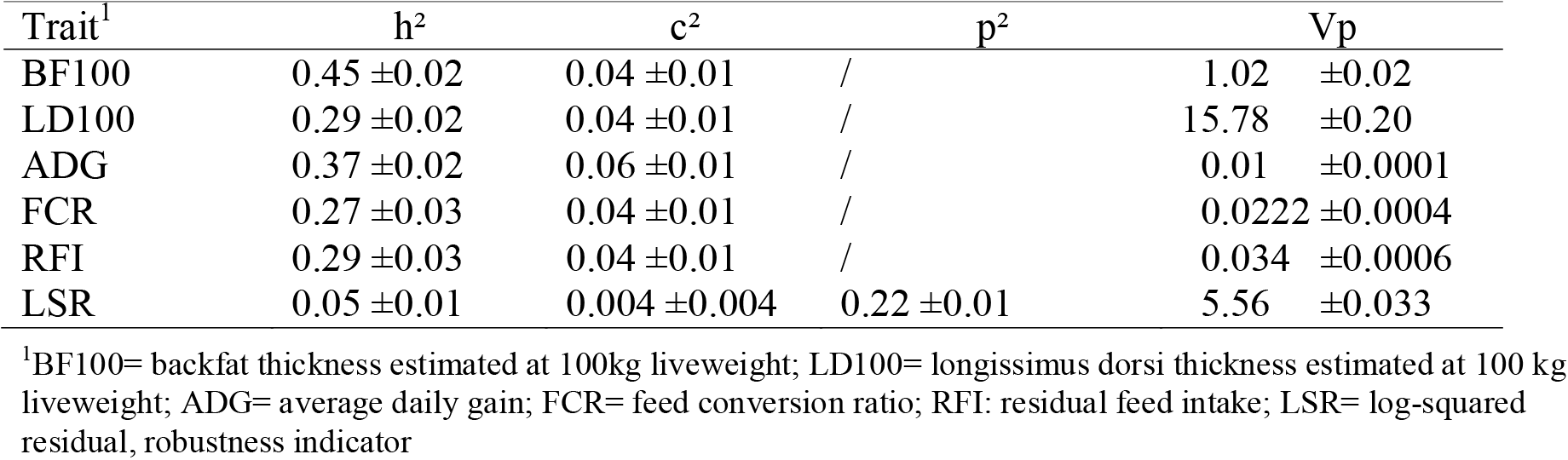
Estimates of heritability (h^2^), common litter effect ratio (c^2^), permanent environmental effect ratio (p^2^) and phenotypic variance (Vp) for the traits recorded (± standard error)

The trait LSR had high negative genetic correlations with ADG, FCR and RFI, ranging from −0.83±0.06 to −0.71±0.06 (Table 3). Estimates of genetic correlations of LSR with BF100 were low and negative, and not significantly different than from 0 with LD100. The trait FCR had a high genetic correlation with RFI, 0.90±0.02, and moderate genetic correlations with ADG and BF100, 0.52±0.06 and 0.50±0.05, respectively. Estimates of the genetic correlations of ADG with BF100 and RFI were positive and moderate to high, 0.43±0.04 and 0.61±0.05 respectively.

**Table 3.**
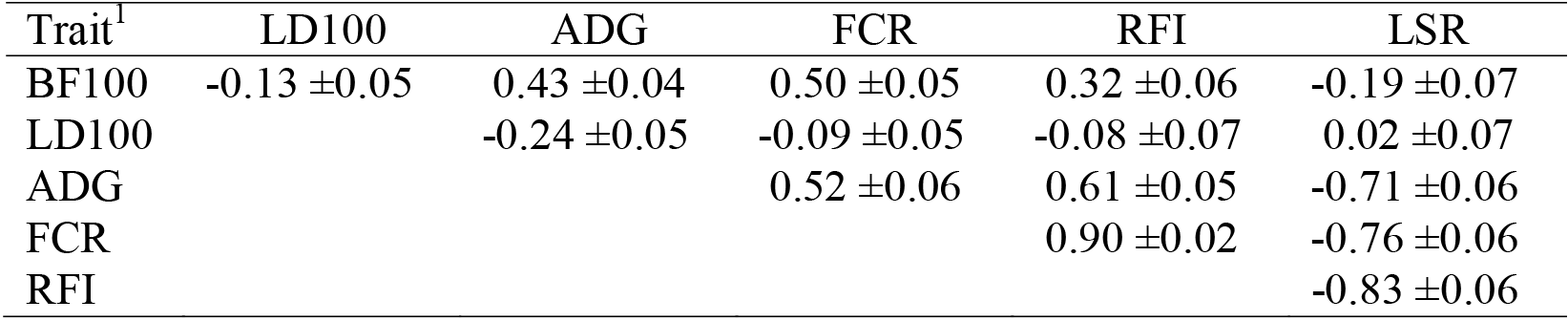
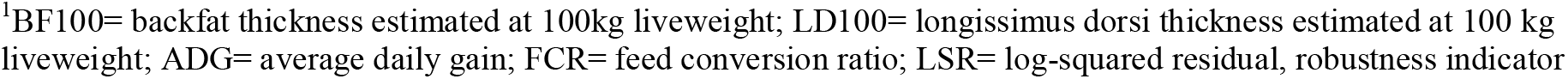
Estimates of genetic correlations (r^2^a ± standard error) between robustness trait (LSR) and production traits

| Trait<sup>1</sup> | LD100 | ADG | FCR | RFI | LSR |
| --- | --- | --- | --- | --- | --- |
| BF100 | -0.13 ± 0.05 | 0.43 ± 0.04 | 0.50 ± 0.05 | 0.32 ± 0.06 | -0.19 ± 0.07 |
| LD100 | | -0.24 ± 0.05 | -0.09 ± 0.05 | -0.08 ± 0.07 | 0.02 ± 0.07 |
| ADG | | | 0.52 ± 0.06 | 0.61 ± 0.05 | -0.71 ± 0.06 |
| FCR | | | | 0.90 ± 0.02 | -0.76 ± 0.06 |
| RFI | | | | | -0.83 ± 0.06 |

<sup>1</sup>BF100= backfat thickness estimated at 100kg liveweight; LD100= longissimus dorsi thickness estimated at 100 kg liveweight; ADG= average daily gain; FCR= feed conversion ratio; LSR= log-squared residual, robustness indicator

### Relation between EBV LSR classes and collected phenotypes

The percentage of “Selectable” animals was significantly related to the EBV LSR quartile (Figure 4). The quartile Q1, including animals with the lowest EBV LSR value, had the highest value with 91.7% of “Selectable” animals, and the quartile Q4 had the lowest percentage, 61.2%. The difference between each quartile were significant. In the quartile Q1, 75% of the animals didn’t receive any antibiotic or anti-inflammatory injection (“No injection”) over the control period. This percentage was not significantly different than those observed for Q2 and Q3, 74.1% and 70.9% respectively. The difference of percentage animals with “No injection” was significant between Q4, 68.7%, and Q1 or Q2. The proportion of animals “Selectable without injection” was significantly higher in Q1 than in Q3 and Q4, 69.3%, 58.3% and 43.3% respectively. In summary, a lower EBV LSR, *i.e*., a higher robustness level, was associated with a better chance of being in good health, of being “selectable” and with a lower use of medicines.

**Figure 4.**
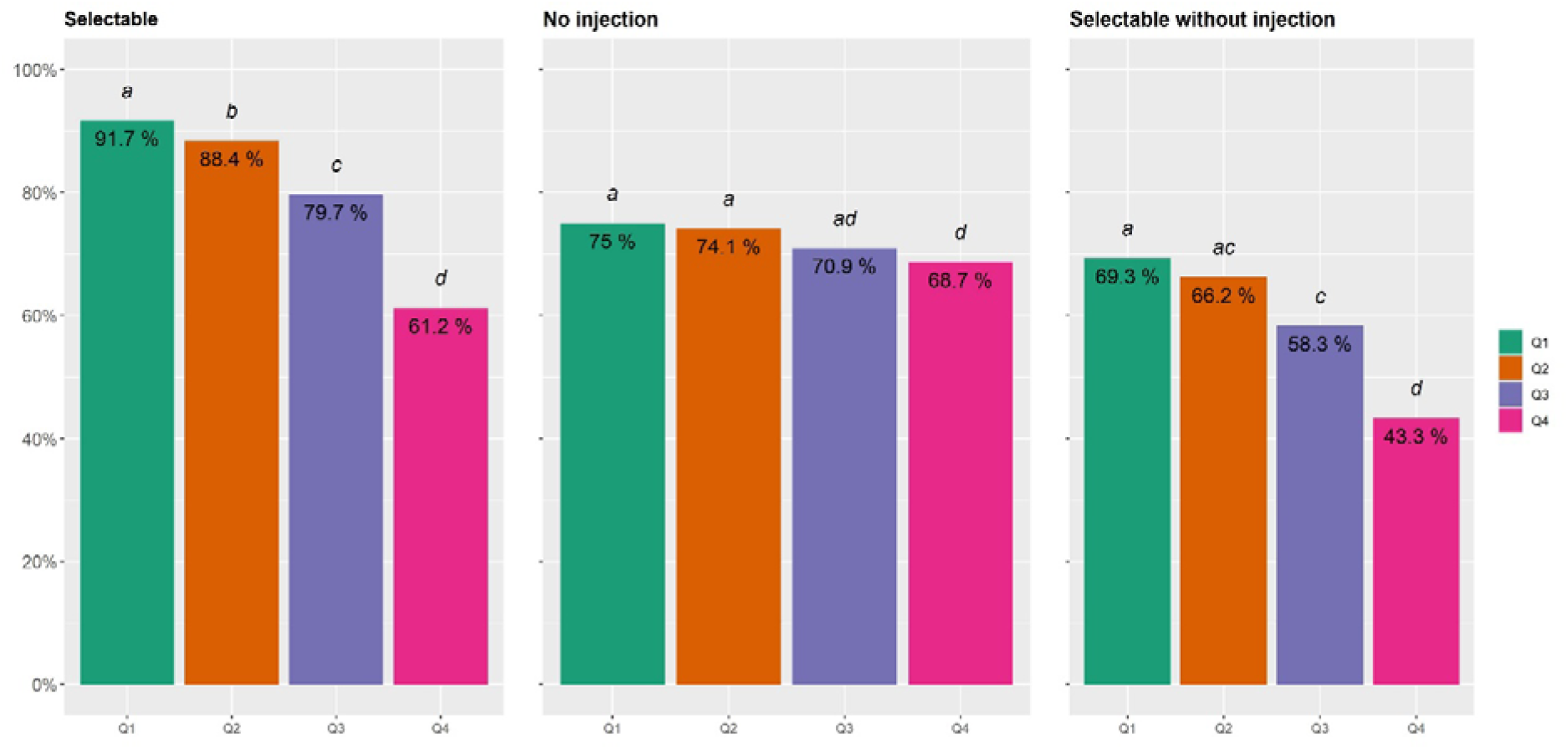
Distribution of percentages of pigs that can be selected (Selectable), that didn’t receive any antibiotic or anti-inflammatory injection (No injection) or that were “Selectable” without receiving any antibiotic or anti-inflammatory injection during the testing period (Selectable without injection) depending of their estimated breeding value for robustness indicator (LSR) quartile. *Q1: pigs with lowest EBV LSR values, i.e., higher robustness genetic potential; Q4: pigs with highest EBV LSR values, i.e., lower robustness genetic potential. Bars with different letters are significantly different (P < 0.05)*.

## Discussion

Our objective was to propose a robustness indicator for fattening pigs from the characterization of the energy allocation of the animal. This indicator is expected to be associated with the ability to cope with different types of environment perturbations encoutered, allowing optimal expresion of production potential. The originality of this work was to use two time-series variables measured in order to model longitudinal an energetic allocation coefficient, α_*t*_, over the fattening period. The LSR trait was estimated as the daily deviation of α between the observed values (*i.e*., calculated with the DLM) and the fitted values estimated by the RR model. Then, we studied the genetic background of LSR in order to assess its potential as selection trait for robustness in fattening pigs. This study indicated that LSR had a low heritability trait and showed strong favorable genetic correlation with growth and unfavorable with FCR and RFI.

### Energetic allocation to growth, from concept to model

When faced with one or more environmental disturbances, we can assume that a fattening pig has two types of responses: a change in feed intake pattern or a modification in energy allocation, that is to say a trade-off. These modifications in feed intake or in allocation patterns can affect or not the body weight gain pattern of the animal. This study focused on the second hypothesis with the objective to quantify robustness with a proxy estimated from variations in the energetic allocation over time. To our knowledge, this approach has been little studied in pigs with a selection purpose. The effects of environmental conditions on feed intake have been widely studied in pigs, mainly the effects of temperature (Quiniou et al., 2000) and diseases (Kyriazakis et al., 1998). The quantification of robustness or resilience through the analysis of variations in feed intake have also been studied (Putz et al., 2019; Nguyen-Ba et al., 2020; Homma et al., 2021). With respect to robustness, the effect of disturbances on growth pattern has been studied on pigs after weaning (Revilla et al., 2019) or during the finishing period (Revilla et al., 2022).

Conceptually, for a fattening pig, it can be assumed that net energy is allocated between several functions: maintenance, growth (daily protein and lipid deposition) and other functions such as health or thermoregulation (Figure 1). We can assume that the proportion of the total available net energy allocated to each function was regulated by a “valve” which increases or decreases allocation over time. This modulation supposed that there would be a regulation in the allocation of the net energy which would be linked on the one hand to a “desired allocation”, dependent on the characteristics of the individual (genotype, age), and on the other hand to an “allocation permitted by the environment”.

The model structure does not detail the full process as described in Figure 1, but provides a simple and biological way to represent energy allocation. Based on these assumptions and on data available in the context of the study, we built the model to estimate α_*t*_ based on daily feed intake and live weight measurements over time. Energy allocation to maintenance was estimated from the metabolic body weight based on the equation proposed by Noblet et al. (1999), although this is an average estimate and we thus ignored any variability between sexes, breeds and individuals. The mobilization of lipid reserves, allowing an increase in the net energy available, was not integrated into the model. Indeed, the mobilization of body reserves, apart from glycogen, is rare in growing animals (van Milgen and Noblet, 2003). In this context, we used a pragmatic approach to estimate the energy available for growth at time *t*. This pragmatic approach is linked to the fact that it is not possible, in a large population, to evaluate precisely for a given pig at a given time, the net energy allocated to maintenance, to additional thermoregulation or physical activity, to protein deposition and to lipid deposition.

In this study, we use DLM regression to model the relation between CNEA_*it-1*_ and CW_*it*_ over time because the DLM makes it possible to characterize allocation coefficient dynamics by a stochastic process, without the requirement for a strong deterministic assumption. With this method, it is possible to determine whether the allocation coefficient was increasing, decreasing or stagnating, without assuming that it followed any given analytical trend, such as a linear, quadratic or cubic trend (Michel and Makowski, 2013). Our approach takes advantage of the available dlm package in R (Petris et al., 2009) which enabled processing of the full data in a small computation time (around 35 min for the 405104 measurements). In addition, this simple DLM approach could ultimately be expanded to the development of multivariate models or the implementation of fixed (batch, herd…) or random effects (Stygar and Kristensen, 2016). Another property of DLM is to produce one-step-ahead forecasts of one or several variables especially to provide early warning to the farmer when forecast error increases (Jensen et al., 2017). Dynamic linear models look to be powerful tools for analyzing time-series variables.

### Estimation of genetic variance in allocation coefficient α_*t*_

We assumed that the “desired allocation” of net energy to growth was driven by two components: the animal’s genetic potential and its degree of maturity. In the first step, the objective was to estimate the genetic variance in allocation coefficient α as affected by degree of maturity, which evolves with the age of the pig. To achieve this, we used a RR model to estimate the genetic variance α_*t*_ and the slope of allocation coefficient to growth over time for each individual. Random regression using orthogonal polynomials models have been widely used in genetics, for example to model feed intake or RFI in pigs or in rabbits (David et al., 2015; Shirali et al., 2017). The random regression of order one was chosen to fit the additive genetic and permanent environmental effects, there was no significant improve of the model, based on LRT test, with polynomials of higher order. If the end of the measurement period had been at a weight closer to the maturity weight, a quadratic random regression would probably be more suitable (Lewis and Emmans, 2020).

The trait α_*t*_, describing the allocation of net energy in growth during fattening period, has moderate heritabilities in the same range as those estimated for FCR or RFI and was strongly correlated with them. In a previous study (Lenoir et al., 2022c), the trait considered was the average value of α_*t*_ and not the repeated estimates, the heritability obtained was lower (0.16±0.05) but was estimated from a different dataset. For the trait RFI, the study of David et al. (2021) showed heritabilities ranging from 0.19±0.06 to 0.28±0.06, using a RR model with weekly estimation over 10 weeks in pigs.

### Genetic parameters for LSR and production traits

The heritability of the trait LSR, which characterizes the environmental variance of α_*t*_, was low but non null. Generally, the heritability of environmental variance is lower than 0.10 (Mulder et al., 2007). This estimate for LSR was in the same range as those published on different traits but with a similar REML method such as: 0.012±0.004 for rabbits birth weight (Garreau et al., 2008), 0.024±0.002 for litter size in pigs (Sell-Kubiak et al., 2022), 0.029±0.003 to 0.047±0.004 for broiler chicken body weight (Mulder et al., 2009). Other studies have been based on the analysis of the log-transformed variance (LnVar) of residuals resulting from a modeling of one time-series variable. This LnVar trait seems to have higher heritabilities than LSR: from 0.20 to 0.24 for milk production (Poppe et al., 2021) or from 0.10 to 0.12 for egg production (Bedere et al., 2022). Some authors have used the double hierarchical generalized linear model (DHGLM) allowing in the same structural model to estimate the mean of the trait and its residual variance (Rönnegård et al., 2010). In order to perform the multi-trait analysis, we chose to use a 2-step approach rather than the DHGLM. In theory, the DHGLM model would make it possible to estimate a residual genetic variance close to the results obtained by our two-step approach. However, it is much more complex mathematically and has convergence issues, making it difficult to use in an operational breeding program (Berghof et al., 2019a).

Heritability estimates for ADG and RFI were consistent with those reported in literature for Pietrain or Large-White pigs raised in similar environmental conditions, which varied from 0.33 ±0.03 to 0.48 ±0.06 and from 0.21 ±0.03 to 0.34 ±0.05 (Saintilan et al., 2013; Déru et al., 2020). For carcass traits (BF100 and LD100), heritabilities were also consistent with the values estimated by Sourdioux et al. (2009) and Saintilan et al. (2013) in the Pietrain breed (BF100: 0.38 to 0.48; LD100: 0.25 to 0.34). Our estimate of heritability for FCR was lower than the heritabilities presented by Saintilan et al. (2013), Gilbert et al. (2017) and Déru et al. (2020), which varied from 0.30 ±0.0 to 0.47 ±0.08.

### Genetic correlations between robustness and production traits

The growth trait ADG was strongly correlated with LSR. In the present rearing conditions, an animal’s ability to be robust, *i.e*., to have low LSR value, is strongly linked to its ability to express optimal growth regardless of the environment. Growth has been a major selection trait in the Pietrain breed for over 20 years, and lack of growth was a major cause of culling at testing or of non-selection. Nonetheless, even if the correlation was strong, it was not equal to 1, which implies that the trait LSR added an additional information regarding the robustness of the animal compared to ADG. Thus, if selection is made using these traits, they would allow us to improve animal’s robustness more than if the selection is made only on growth traits.

There were strong and unfavorable relationship between LSR and feed efficiency traits, FCR and RFI. This could be related to the positive correlation between ADG and FCR, which was affected by the way these two traits were estimated (Lenoir et al., 2022a). The traits ADG and FCR used in selection were measured over an identical period for all pigs but were not standardized between starting and finishing weights. Accordingly, some of the animals tested reached their mature weight before testing, which led to a drop in feed conversion or residual feed intake even if they had previously a strong growth. Thus, there were two different types of finisher pigs with low FCR or RFI: those which had a strong growth but did not approach their mature weight during the testing period, and those with a low daily feed intake associated with a low, near maturity, growth (Lenoir et al., 2022a). We performed an additional analysis where we standardized the trait FCR between 40 and 100kg, the genetic correlation with LSR remained unfavorable but less strong, −0.34±0.14. The standardization of the trait FCR modified the genetic correlation with ADG from moderately unfavorable, 0.52±0.06, to close to zero or slightly favorable, −0.08±0.09. The correlation between LSR and FCR or RFI could indicate that the most robust pigs during the testing period were not the most efficient because they allocate a part of energy to other functions or maintenance. Indeed, a selection for low RFI could impact the ability of the animals to modify their allocation of energy to other functions to cope with environmental challenges (Gilbert et al., 2017). This antagonism between short-term efficiency and resilience has been put forward by Friggens et al. (2017). In this situation, it would seem that there is a compromise that does not make it possible to increase robustness relatively easily without loss of selection response in feed efficiency. In contrast, several studies have shown through divergent selection experiments on RFI, that animals from Low RFI line (LRFI) adapted better to environmental challenges or at least are not disadvantaged compared to animals from High RFI line (HRFI). Chatelet et al. (2018) showed that the health, growth performance and feed intake of animals from the LRFI line were less impacted than those of animals born from the HRFI line under poor hygienic conditions. In the same selection experiment, the risk of being culled between 70 days of age and slaughter was 1.8 times less in the LRFI line compared to the HRFI line (Gilbert et al., 2017). In another experience of selection Dunkelberger et al. (2015) suggested that pigs for LRFI were more robust to PRRSV challenges; their growth was less affected and they was less affected. These results seem to contradict the resource allocation theory and the genetic correlation estimated in our study. This study was carried out on Pie NN line, a sire line, and the different selection experiments on RFI were realized with animals from Large-White (or Yorkshire), a dam line. The Pietrain sire line had been created and selected for several generations on objectives of improving feed efficiency, growth and carcass characteristics, potentially to the detriment of the other traits, such as robustness. Due to these characteristics and orientations, it can be assumed that there is a different allocation pattern between these lines.

Genetic correlations between robustness and BF100 were slightly unfavorable. We can suppose that the capacity to be robust could be associated with more important body reserves allowing the animal to face perturbations.

### Relation between EBV LSR classes and collected phenotypes

Our study shows that model longitudinal energetic allocation to growth offers the opportunity to develop a proxy of robustness that is heritable. Further, this proxy has to meet the expectations of pig farmers, that is to say, it should identify animals that faced to environmental disturbances and were present for testing in good health and with the least amount of medical injections. The analysis of the relationship between EBV LSR quartiles and phenotypes routinely collected on farms shows the most favorable situation for the most robust animals, *i.e*., those from the quartile with the weakest EBV LSR (Q1). Including LSR in the breeding goal would be an opportunity to improve the robustness qualities of Pie NN line for the fattening period, in spite of the low heritability of LSR. However, these results are evaluated over a short period of animal’s life, it would be appropriate to investigate the effects of a selection on the LSR trait over the whole lifespan of related animals (dam, sire, pure or crossbred offspring). In a following step, it could be relevant to study the link between the LSR trait and the reproductive performances of boars (spermatic production) or females (fertility, productive longevity, survival).

### Environmental conditions

This studied was carried out in a higher biosecurity environment than regular farms, related to the fact that a breeding company cannot take any risks with a purebred nucleus. The other environmental conditions (feed characteristics, barn design, density…) were close to those found in production farms in France, that is, designed to minimize exposure to environmental challenges. When designing the selection conditions there is a need to balance between conditions that allow full expression of performance and meet sanitary requirements versus conditions that favor expression of robustness. Even though these environments are qualified as favorable, the animals are subjected to stresses which can be chronic (social stress, heat wave). Rear animals under challenging conditions seems to allow better phenotyping of the robustness of the animals (Gunia et al., 2018). The difficulty of having conditions to evaluate robustness while evaluating production potential could be partly overcome by the use of short-term challenges, such as feeding challenges. Indeed, offspring of these purebred pigs, selected in one type of environment, are likely to be reared in harder and more variable environments impacting robustness expression. This relationship between the robustness expression and diverse rearing conditions cannot be dissociated from Genotype x Environment (GxE) interaction (Falconer and Mackay, 1996). This interaction that may cause reranking of sires, has a greater impact on traits based on variances than on traits based on means (Bedere et al., 2022). The acquisition of data on related animals reared in farms newly equipped with AFS makes it possible to consider evaluating the effects of the GxE interaction.

In this study, we proposed an approach for characterizing the robustness through the variability in the allocation. However, when studying the allocation pattern, it is important to also assess the acquisition trajectory (van Noordwijk and de Jong, 1986; Friggens et al., 2017). In a routine selection approach, it would be relevant to add to the LSR trait, a trait making it possible to characterize the robustness on acquisition

## Conclusions

The trait LSR could be interpreted as an indicator of the response of the animal to perturbations/stress, that is to say a robustness proxy. This study shows that LSR has a low heritability but that it is possible to set up a selection on this trait. We found that this trait is favourably genetically correlated with a growth trait (ADG) and unfavourably with feed efficiency traits (FCR and RFI). Estimation of the economic value of LSR trait is a key issue before adding this trait in breeding goals. Furthermore, improving robustness qualities also meets societal expectations, the economic value of which is difficult to quantify.

## List of abbreviations

α_*t*_: daily energetic allocation coefficient to growth
ADG: average daily growth
AFS: automatic feeding system
AMW: average metabolic weight
BF: backfat thickness
BF100: backfat thickness estimated at 100 kg liveweight
BW: body weight
CNEA: cumulative net energy available for growth
CW: cumulative weight gain
DFI: daily feed intake
DLM: dynamic linear model
EBV: estimated breeding value
EI: net energy intake
FCR: feed conversion ratio
FI: feed intake
IBW: initial body weight
LD: longissimus dorsi thickness
LD100: longissimus dorsi thickness estimated at 100 kg liveweight
LSR: log transformed squared residuals, robustness indicator
MR: net energy maintenance requirement
NEA: net energy available for growth
PDFI: potential average daily feed intake
Pie NN: Piétrain NN Français free from halothane-sensitivity
RFI: residual feed intake
TBW: body weight at individual testing

## Declarations

### Ethics approval and consent to participate

Specific Experimental Animal Care and Use Committee approval was not needed because all the data used in this study were obtained from preexisting databases provided by AXIOM. The data used were from animals raised under commercial conditions that were cared for according to EU-Council directive 2008/120/EC of 18 December 2008 laying down minimum standards for the protection of pigs (http://data.europa.eu/eli/dir/2008/120/oj).

### Consent for publication

Not applicable

### Availability of data and materials

The datasets analysed during the current study are not publicly available because they are part of the commercial breeding program of AXIOM. However, they are available from the corresponding author on reasonable request.

## Competing interests

RMT, NF and ID declare that they have no competing interests. GL and LFG are employed by AXIOM. The datasets are of interest to commercial targets of AXIOM, but this interest did not influence the results in this manuscript in any matter.

### Funding

This project was supported by ANRT (Association Nationale Recherche Technologie) with a CIFRE Doctoral fellowship (2019/0705).

## Authors’ contributions

GL, NF and RMT developed the conceptual model. RMT and GL implemented and tested the DLM model. GL, ID and LFG carried out statistical analyses. GL drafted the paper. ID, LFG, NF, RMT and GL participated in interpreting and discussing results. All authors read and approved the final manuscript.

## Acknowledgements

The authors would like to acknowledge the breeding company Axiom for funding GL’s Ph.D thesis. We also acknowledge the technicians of AXIOM’s testing station at La Garenne for their implication for data collection. We also thank Céline Lebeux and Kambiz Kashefifard for managing data collection on farm and development of the database to store and share the data.

## Additional information

**Appendix 1.** List of individuals observations performed during the individual test from Lenoir et al. (2022a).

|  | Observation |
| --- | --- |
| Observations taken into account to<br>define the robustness traits | Abcess |
|  | Cannibalism |
|  | Capelet |
|  | Weak development / Low body condition |
|  | Callus |
|  | Shortness of breath |
|  | Necrotic ear |
|  | Out of test (testing body weight < 70kg) |
|  | Shaker |
| Observations not taken into account<br>to define the robustness traits | Lack of leg soundness |
|  | Low and short |
|  | Conformation / Body development |
|  | Culard |
|  | Important conformation |
|  | Fat animal |
|  | Asymmetric hooves |
|  | Teats default |
|  | Incorrect conformation |
|  | Hernia |

## Notes

### Summary of Updates

Modification of authors affiliations

